# Proliferative cell targeting and epithelial cell turnover fuels hepatitis E virus replication in human intestinal enteroids

**DOI:** 10.1101/2024.03.27.586953

**Authors:** Nanci Santos-Ferreira, Xin Zhang, Laura Corneillie, Jana Van Dycke, Winston Chiu, Claire Montpellier, Johan Neyts, Laurence Cocquerel, Suzanne J. F. Kaptein, Joana Rocha-Pereira

**Affiliations:** KU Leuven, Department of Microbiology, Immunology and Transplantation, Rega Institute, Virus-Host Interactions & Therapeutic Approaches (VITA) Research Group, Leuven, Belgium; KU Leuven, Department of Microbiology, Immunology and Transplantation, Rega Institute, Virology, Antiviral Drug & Vaccine Research Group, Leuven, Belgium; Univ. Lille, CNRS, Inserm, CHU Lille, Institut Pasteur de Lille, U1019-UMR 9017-CIIL-Center for Infection and Immunity of Lille, F-59000 Lille, France

**Keywords:** Hepatitis E virus, Human intestinal enteroids, Infection, Stem cells, Enteroendocrine cells, Proliferating cells

## Abstract

Hepatitis E virus (HEV) is a leading pathogen causing acute viral hepatitis globally. While HEV is primarily spread fecal-orally, the role of the gut in HEV pathogenesis remains largely unexplored, including how HEV disseminates from gut to liver, and whether the gut is an HEV reservoir. We here aimed to illuminate HEV biology in the gut using human intestinal enteroids (HIEs).

Three strategies were explored to establish an HEV-HIE model: three-dimensional (3D) HIEs, two-dimensional (2D) HIEs in transwell, and HEV RNA-electroporated HIEs. HEV particles produced by electroporated HIEs were characterized by western blot and gradient centrifugation. The intestinal tropism of HEV was investigated through confocal fluorescent microscopy and gene expression analysis.

HEV infection in 3D-HIEs and 2D-HIEs showed limited replication, whereas HIEs electroporation led to a sustained increase in the release of non-enveloped infectious virions. These virions could re-infect new 3D-HIEs, yielding a ∼2 log_10_ increase in HEV RNA. In electroporated HIEs, high expression of the infectious ORF2 capsid form was observed in the supernatant. Importantly, 70% of all HEV-infected cells were identified as proliferative cells. ORF2 staining was also observed in absorptive enterocytes, goblet, and enteroendocrine cells.

Overall, we established a robust HEV-HIE model that yields high titers of infectious non-enveloped virions. Proliferative cells and the fast intestinal epithelial cell turnover are important features that facilitate efficient HEV replication, and likely also its dissemination. This study suggests that the gut is an HEV reservoir, capable of producing some of the non-enveloped HEV shed in the feces.

**Significance:** Hepatitis E virus (HEV) causes ∼20 million enterically transmitted cases of viral hepatitis globally. To better understand the largely unexplored role of the gut during an HEV infection, we used human intestinal enteroids (HIEs). We show that HEV-electroporated HIEs produce highly infectious non-envelopedHEV particles, resulting in a more efficient re-infection compared with traditional gold-standard methods. Importantly, we detected HEV in multiple human intestinal epithelial cell types within the human intestinal epithelium, with the majority of proliferative cells infected. This model offers valuable insights into the intestinal tropism of HEV and can serve to further investigate chronic HEV infection in the gut. Moreover, this underscores the importance of using physiologically relevant models to unravel details of HEV infection in the gut.

## Introduction

The enteric hepatitis E virus (HEV) poses a substantial global health burden. HEV is mainly transmitted fecal-orally, with human-infecting genotypes predominantly belonging to the *Paslahepevirus balayani* species (*Hepeviridae* family)^1^. HEV-1 and - 2 are endemic in developing regions with an estimated 70,000 deaths annually^2,3^. These infections occasionally result in fulminant hepatitis in pregnant women, with mortality rates reaching 20 to 30%^4^. Zoonotic HEV-3 and -4 cause sporadic autochthonous infections in developing and industrialized regions^2^. HEV-3 also causes chronic infections, especially in immunocompromised individuals, thereby increasing the risk of rapid progression to cirrhosis, graft loss and death^5^. There is no approved specific antiviral therapy. The off-label use of ribavirin for chronic infection is associated with marked adverse effects and treatment failure^6–8^.

HEV presents in two forms in infected hosts: quasi-enveloped (HEVenv^+^) and naked/non-enveloped particles (HEVenv^-^)^9^. During infection, HEVenv^+^ is released from the apical membrane into the bile duct, where the lipid envelope is presumably degraded by detergents and proteases in the bile, explaining the appearance of HEVenv^-^ in feces and bile^10^. While HEVenv^+^ circulates in the blood, HEVenv^-^ is presumed to be the main form responsible for transmission and is associated with higher infectivity^11,12^. In standard cell culture systems, intracellular HEV predominantly exists as HEVenv^-^, while HEVenv^+^ is secreted into the supernatant^13^. The HEV genome encodes three partially overlapping open reading frames (ORFs)^14^. ORF2 encodes the major structural (*i.e.,* capsid) protein that can manifest in three different forms: ORF2 is either assembled into infectious particles (referred to as ORF2i) and secreted via the exosomal pathway from infected hepatocytes, or secreted in glycosylated (ORF2g) and cleaved (ORF2c) forms not associated with infectious particles^15^.

Numerous knowledge gaps remain on fundamental aspects of HEV infection and pathogenesis, partly due to challenges in HEV cultivation (*i.e.,* slow replication to low titers)^16^. Since HEV is hepatotropic, the virus is typically propagated *in vitro* using liver-derived primary cells and hepatic immortalized cell lines. Despite HEV also being an enteric virus that is transmitted fecal-orally and shed in high titers in stool^9^, the role and contribution of the gut in HEV-induced disease has received limited attention. HEV was shown to replicate in intestinal epithelial cells that release infectious particles, particularly from the apical side, which ribavirin failed to block, possibly explaining viral relapses observed in chronically infected patients^17^. The gut epithelium was proposed to serve as the initial virus amplification site before dissemination to the liver, and acting as a reservoir, especially during chronic infections^17,18^. Yet, how the virus migrates from the gut to the liver is still not well understood. Moreover, a detailed understanding of HEV infection in the gut is still lacking, particularly concerning the specific intestinal cell types that are infected, the host response to the infection, the viral dissemination routes to extra-intestinal tissues (*e.g.,* kidney and central nervous system), and the disease-inducing mechanisms. Thus, new and more physiologically relevant cultivation models are needed to recapitulate the full tropism of HEV and illuminate missing details in HEV biology.

Organoid technology has the potential to advance knowledge in virus biology and host-pathogen interactions by providing a more physiologically relevant model. Notably, HEV replication in human liver-derived organoids has been reported^19^. Human intestinal organoids (HIOs) closely mimic the human intestinal epithelium *in vitro*, making them valuable for dissecting virus-host interactions of several enteric viruses^20–25^. HIOs are non-transformed three-dimensional (3D) cell cultures arranged in a crypt-villus structure that incorporate the physiological features of the intestinal epithelium, including the presence of different cell populations (enterocytes, goblet cells, enteroendocrine and Paneth cells)^26^. Intestinal organoids can be derived from induced pluripotent stem cells or tissue containing Lgr5+ (leucine-rich-repeat-containing G-protein-coupled receptor 5) stem cells, known as human intestinal enteroids (HIEs)^27,28^. We here used HIEs to establish an HEV infection model of the small intestine via various strategies and subsequently employed it to study HEV infection, including its cellular tropism in the gut^29^.

## Results

### HEV infection in differentiated 3D human intestinal enteroids is limited

To investigate whether HIEs are permissive to HEV replication, fetal and adult differentiated 3D-HIEs were infected with wildtype HEV-3 Kernow-C1 p6 or Kernow-C1 p6 G1634R bearing a fitness-enhancing mutation^6^ (for simplicity, we will below refer to these as respectively ‘HEV-3’ and ‘HEV-3^G1634R^’), after which the viral load (derived from pooled cell extracts and supernatant) was assessed at different days post-infection (pi) (Fig. 1A). When using an inoculum of 1.0×10^7^ HEV RNA copies, HEV-3 infection of either fetal or adult HIEs resulted in a very limited gain in HEV RNA over input (Fig. 1B). Infection with higher titer HEV-3^G1634R^ (5.0×10^7^ RNA copies) yielded a modest increase of ∼1 log_10_ in HEV RNA from 6 to 72 hours pi, after which levels remained stable until day 5 pi, albeit significant in adult HIEs (Fig. 1C). Thus, a high inoculum may be required to kick-start the infection but results in no gain in replication.

**Fig. 1.**
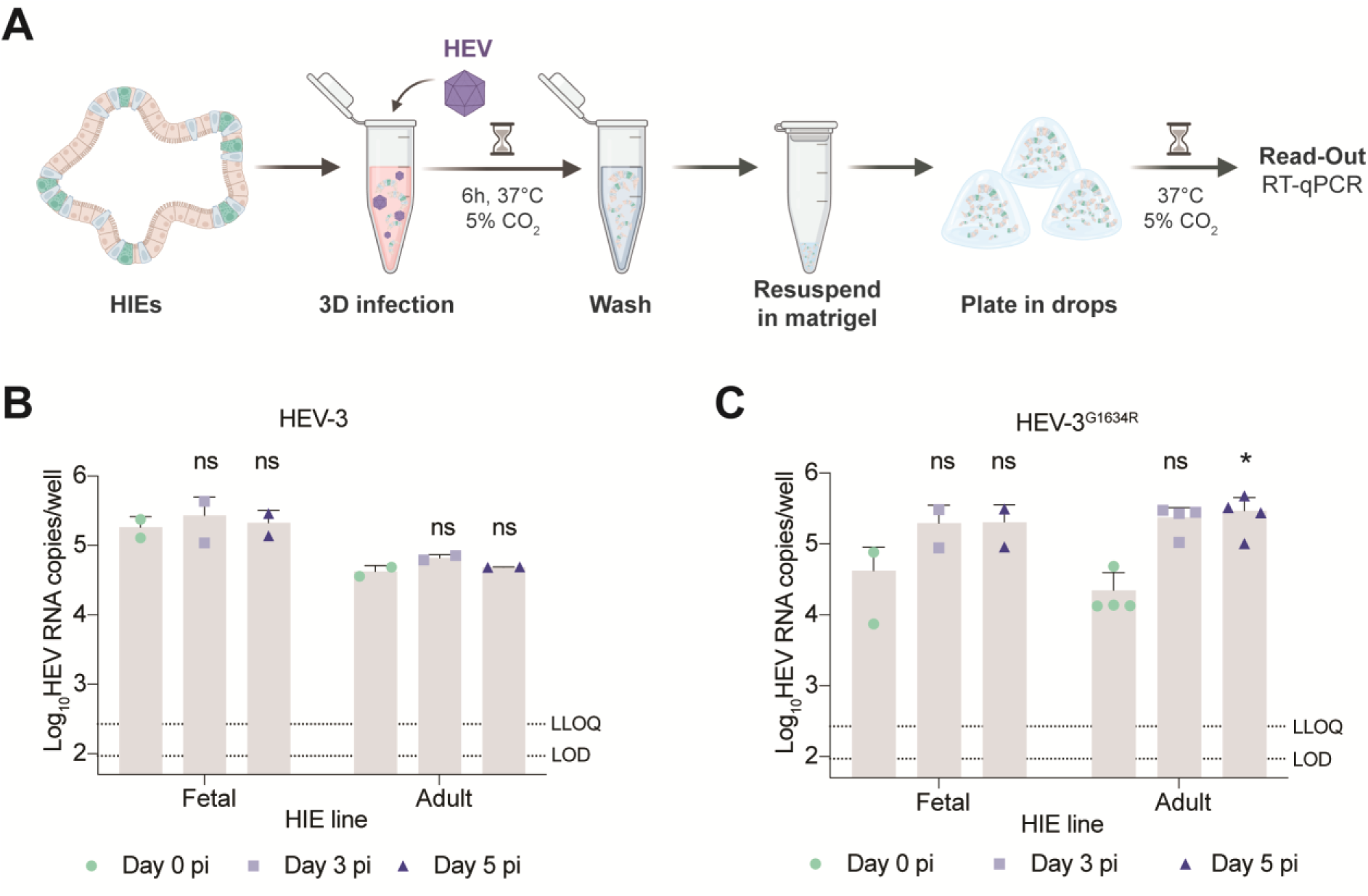
Establishment of HEV infection in the 3D-HIEs model. (A) Schematic representation of the experimental layout. (B) Fetal and adult differentiated 3D-HIEs were infected with HEV-3 (env^-^, 1.0×10^7^ GEs) (*N*=2). (C) Fetal (*N*=2) or adult (*N*=4) differentiated 3D-HIEs were infected with HEV-3^G1634R^ (env^-^, 5.0×10^7^ GEs). HEV RNA of the whole well (pool of supernatant and cell lysate) was quantified by RT-qPCR. Day 0 pi represents 6 h pi. Data are mean+SD. *, *P*≤0.05. Statistical analysis was performed using two-way ANOVA followed by Tukey’s test for multiple comparisons. GEs, genome equivalents; LLOQ, lower limit of quantification; LOD, limit of detection; pi, post-infection. Schematic representation created with BioRender.com.

### HEV infection in HIE polarized monolayers leads to higher apical viral shedding

Next, we cultured HIEs as polarized monolayers (2D) in a transwell system, allowing us to study virus shedding into both the apical and basolateral compartments (Fig. 2A). Prior to HEV infection, the transepithelial electrical resistance (TEER) of the monolayers was measured at >300 Ω/cm^2^, indicating a tight and polarized intestinal epithelium. Infection of differentiated fetal or adult 2D-HIEs with the (slightly better replicating) HEV-3^G1634R^ led to a constant virus shedding (HEV RNA levels of ∼5.5 log_10_/mL) to the apical side of the intestinal epithelial layer (Fig. 2B). Basolateral shedding was also detected, albeit at lower levels of ∼3.5 - 4 log_10_ HEV RNA copies/mL (Fig. 2B). Given that half of the culture media was replaced by fresh media in both compartments every two days, this result demonstrates that new virions were produced and shed into the culture supernatant. Intracellular HEV RNA levels remained around 5 log_10_ copies/well with a slight drop on day 5 pi (Fig. 2C).

**Fig. 2.**
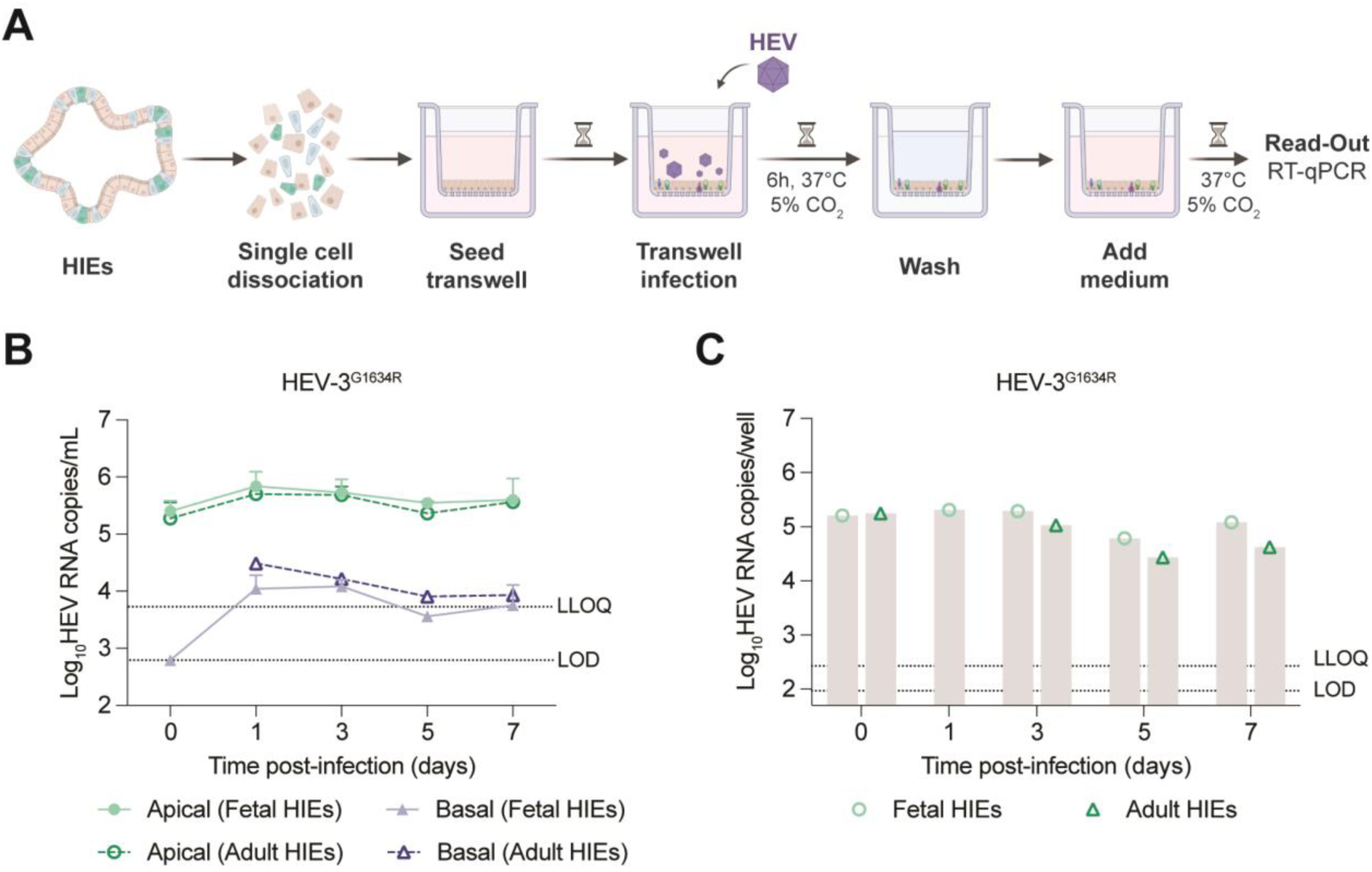
HEV infection dynamics in polarized 2D-HIEs in transwell system. (A) Schematic representation of the experimental layout. (B) Extracellular HEV released into apical or basal supernatant of fetal (*N=2*) or adult (*N=1*) differentiated 2D-HIEs infected with HEV-3^G1634R^ (env^-^, 2.0×10^8^ GEs). Half of the medium (100 µL apical and 300 µL basolateral compartment) was refreshed every 2 days. (C) HEV RNA levels in infected fetal or adult differentiated 2D-HIEs cell lysates. Day 0 pi represents 6 h pi. Data are mean+SD.

### Robust HEV replication in HIEs electroporated with capped viral RNA

Next, we evaluated whether electroporation of a HIE single-cell suspension with capped viral RNA resulted in a more efficient HEV replication. Single cell HIEs were electroporated with subgenomic or full-length HEV capped RNA from either human HEV-1 (Sar55/S17), HEV-3 (Kernow-C1 p6), HEV-3^G1634R^ (Kernow-C1 p6 G1634R), or rat HEV (pLA-B350), as illustrated in Fig. 3A. HIEs electroporated with RNA of the luciferase-encoding subgenomic replicons HEV-3/luc, HEV-1/luc or rat HEV/luc yielded a 17-, 5- and 10-fold increase in luciferase levels, respectively, at the day of peak replication (day 4 post-electroporation [pe] for HEV-3/luc and rat HEV/luc, day 11 pe for HEV-1/luc) (Fig. 3B). Ribavirin treatment resulted in lower luciferase signals compared to non-treated HIEs electroporated with HEV-3/luc, with a mean half-effective concentration of around 25 µM (Fig. 3C).

**Fig. 3.**
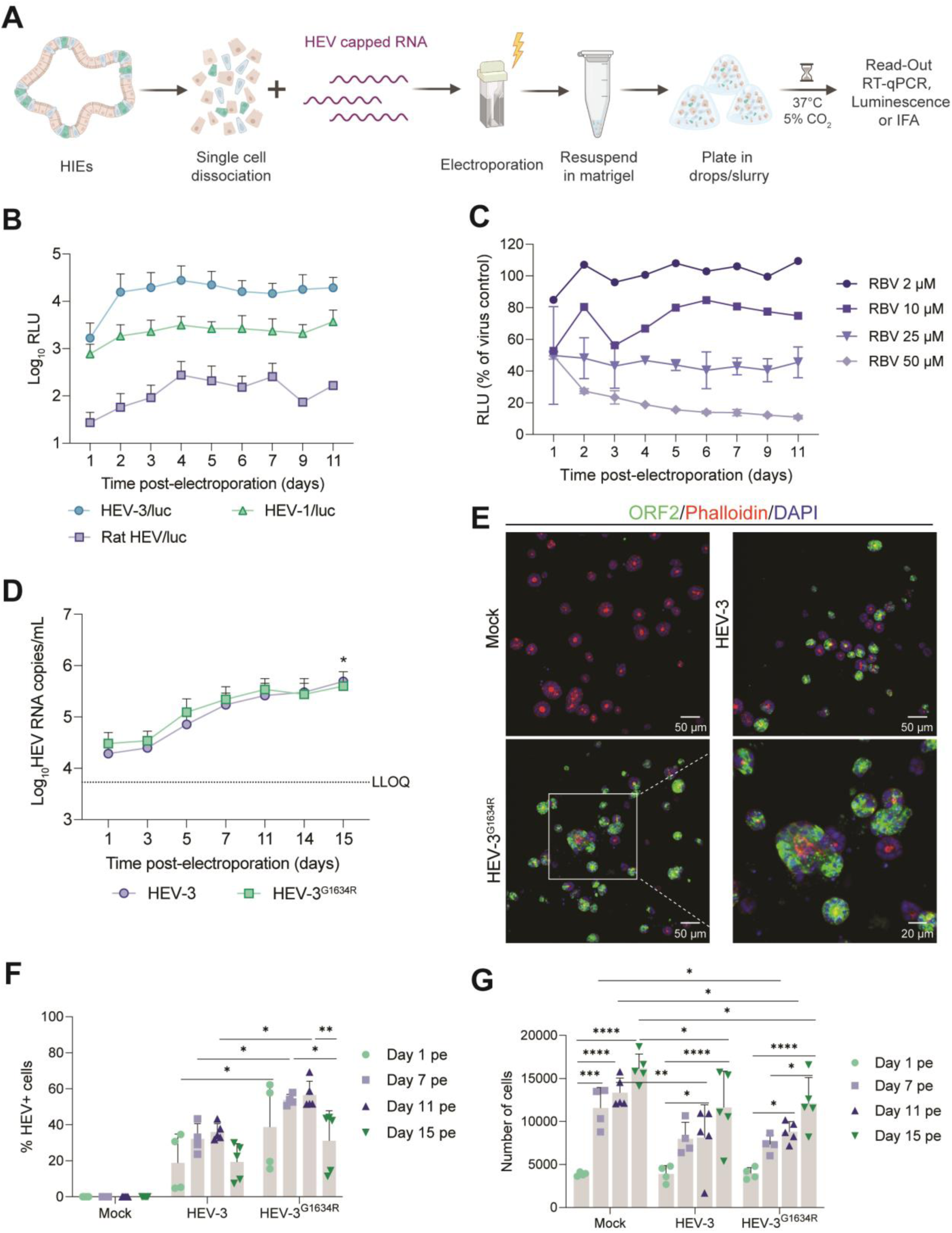
Electroporated HIEs allow robust HEV replication. (A) Schematic representation of the experimental layout. (B) Fetal HIEs were electroporated with mock, HEV-3/luc (*N=6*), HEV-1 HEV-1/luc (*N=2*) or ratHEV/luc (*N=3*) capped RNA. (C) HEV-3/luc electroporated fetal HIEs treated with different RBV concentrations (*N=2*). Viral replication related luciferase activity was determined in 20 µL supernatant up to day 11 pe. (D) Fetal HIEs were electroporated with mock, HEV-3 (*N=4*) or HEV-3^G1634R^ (*N=4*) full-length capped RNA. Half of the medium was refreshed at day 11 pe. Viral replication was determined by RT-qPCR in 50 µL supernatant up to day 15 pe. \**P*≤0.05 (calculated using two-way ANOVA followed by Tukey’s test for multiple comparisons; comparing day 1 pe versus day 15 pe for either HEV-3 or HEV-3^G1634R^). (E) Representative images of HEV ORF2 expression in HEV-3 or HEV-3^G1634R^ electroporated fetal HIEs at day 11 pe. Immunofluorescence staining with ORF2 (HEV, green); Phalloidin (actin, red); and DAPI (nuclei, blue). Images were acquired on a Spinning-disk confocal microscope with a 25× objective. Scale bar – 50 or 20 µm. (F) Percentage of HEV ORF2 positive cells in mock-, HEV-3- or HEV-3^G1634R^- electroporated HIEs at day 1, 7, 11 and 15 pe. The percentage of infected cells are defined as total number of cells containing ORF2 signal within the nucleous or in close proximity thereof divided by the number of total cells counted using the DAPI signals in the same well. (G) Number of cells in mock-, HEV-3 or HEV-3^G1634R^- electroporated HIEs at day 1, 7, 11 and 15 pe. Number of cells was determined by nuclear staining (DAPI) and enumerated by high-content imaging (HCI). Statistical analysis was performed using the 2-way ANOVA, followed by Tukey‘s multiple comparisons test. *, *P*⩽0.05; **, *P*⩽0.01; ***, *P*⩽0.001; ****, *P*⩽0.0001. pe, post-electroporation; RLU, relative luminescence unit; RBV, ribavirin;. Data are mean+/±SD.

Electroporation with full-length HEV-3 or HEV-3^G1634R^ resulted in a sustained increase of the viral load in the supernatant over time, reaching 4.9×10^5^ and 5.8×10^5^ HEV RNA copies/mL, respectively, at day 15 pe (Fig. 3D). Moreover, HIEs electroporated with HEV-3 or HEV-3^G1634R^ markedly expressed ORF2 antigens (Fig. 3E). The percentage of infected cells was 36% and 57%, respectively, in HEV-3 and HEV-3^G1634R^ electroporated HIEs, with the peak replication at day 11 pe (Fig. 3F and S1A-F), in line with viral RNA levels (Fig. 3D). The single-cell electroporated HIEs proliferated and formed the characteristic 3D organoids, translating in a significant increase in the cell number over time (Fig. 3G). HIEs electroporated with HEV RNA, however, grew slower and the 3D structures were smaller compared to their mock-electroporated equivalents (Fig. S1D and G). This difference is likely not due to cell death, as HEV-electroporated HIEs presented less propidium iodide positive staining than did mock-electroporated HIEs (Fig. S2). Ribavirin treatment of full-length HEV-3^G1634R^-electroporated HIEs resulted in a dose-dependent reduction in intracellular and extracellular HEV RNA levels (Fig. S3), corroborating the earlier findings obtained with HIEs electroporated with HEV subgenomic RNA (Fig. 3C).

### HEV infects and preserves the proliferative intestinal stem cell niche

Next, we looked at the cellular composition of the HEV-electroporated HIEs and the permissiveness of various intestinal cell types to HEV infection. Electroporated HIEs were mainly composed of proliferating cells (Ki67+ cells), reaching 45% and 70% of the cell composition, in mock and HEV-electroporated conditions, respectively, in the first week (Fig. 4A). Interestingly, the amount of proliferating cells decreased in mock-electroporated HIEs while it increased in HEV-electroporated HIEs during the first week after electroporation. When looking at the mRNA transcript levels of the intestinal epithelial cell markers, Lgr5+ (stem cells), lysozyme (Paneth cells), sucrose isomaltase (mature enterocytes), mucin 2 (goblet cells) and chromogranin A (enteroendocrine cells), a similar increase in transcript levels was found in HIEs electroporated with HEV-3 and HEV-3^G1634R^ as well as for mock-electroporated HIEs (Fig. S4), suggesting that HEV infection had no effect on the proliferative cell niche, while enabling differentiation of mature cell types, *i.e.,* enterocytes, goblet and enteroendocrine cells.

**Fig. 4.**
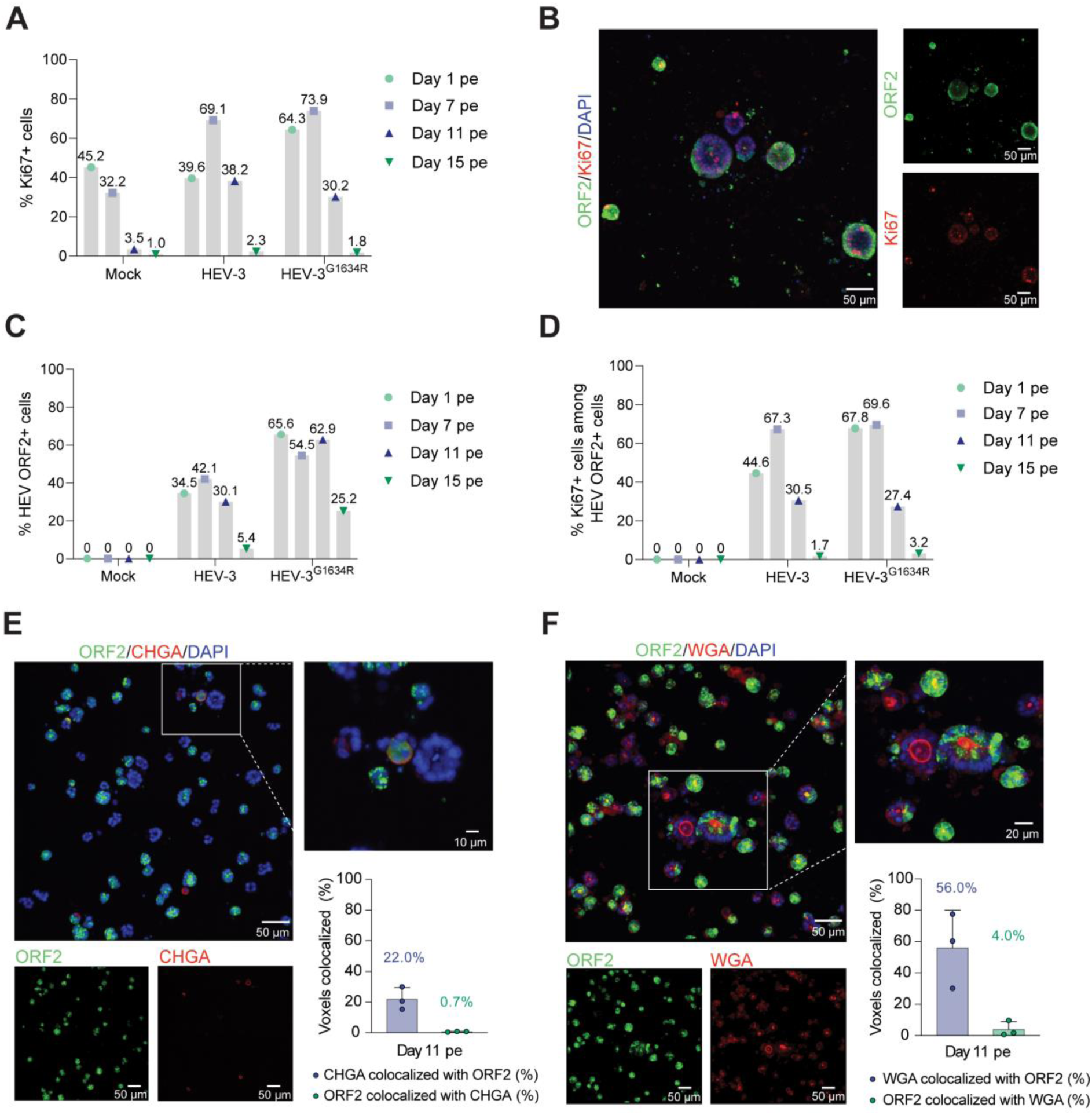
Targeted cell tropism of HEV in infected HIEs. (A) Percentage of proliferative cells in mock-, HEV-3 or HEV-3^G1634R^- electroporated HIEs at day 1, 7, 11 and 15 pe. Amount of proliferative cells was determined by HCI and defined by the objects positive for both DAPI and the proliferation marker Ki67. (B) Representative image of immunofluorescence staining of HEV-3^G1634R^ electroporated fetal HIEs at day 11 pe. Cells were stained with HEV ORF2 (green) and Ki67 (red). (C) Percentage of HEV ORF2 capsid positive cells within the proliferative cell population in mock-, HEV-3 or HEV-3^G1634R^- electroporated HIEs at day 1, 7, 11 and 15 pe. These cells were defined by the objects positive for all three markers (Ki67 and HEV ORF2). (D) Percentage of cells that were positive for proliferative marker (Ki67) within the HEV ORF2 positive population in mock-, HEV-3 or HEV-3^G1634R^ electroporated HIEs at day 1, 7, 11 and 15 pe. These cells were defined by the objects positive for all three markers (DAPI, Ki67 and HEV ORF2). (E, F) Representative images of immunofluorescence staining of HEV-3^G1634R^ electroporated HIEs at day 11 pe. Cells were stained with HEV ORF2 (green) and antibody specific for (E) enteroendocrine cells (CHGA, chromogranin A, red) or (F) goblet cells (WGA, wheat germ agglutinin). DAPI (blue) was used as counterstaining. Overlap of voxels between HEV ORF2 and (E) CHGA signal (*N=3*, 8 fields/experiment) or (F) WGA signal) (*N=3*, 5 fields/experiment). pe, post-electroporation.

Confocal imaging of immunofluorescent staining of virus and cell markers showed that proliferating cells, *i.e.,* positive for the proliferation marker Ki67, were positive for HEV ORF2 protein expression (Fig. 4B, S5 and S6A) and with up to 42% and 55% of the proliferating cells found to be infected with HEV-3 or HEV-3^G1634R^, respectively, on day 7 pe (Fig. 4C). On all HEV-infected cells, almost 70% were proliferating cells (Fig. 4D).

We also examined what other intestinal epithelial cell types were infected, namely mature cell types. HEV ORF2 signal was detected in enteroendocrine cells (chromogranin A, CHGA) (Fig. 4E and S6B). Colocalization of virus with CHGA showed that 0.7% of the thresholded HEV ORF2 signal colocalized with the CHGA signal, while 22.0% of the thresholded CHGA signal overlapped with the ORF2 signal (Pearson coefficient of 0.092). Moreover, ORF2 expression was also detected in cells containing mucins (mucin staining with wheat germ agglutinin, WGA), indicating that HEV is able to infect goblet cells (Fig. 4F). Colocalization analysis showed that 4.0% of the thresholded HEV ORF2 signal colocalized with WGA signal, while 56.0% of the thresholded WGA signal overlapped with the ORF2 signal (Pearson coefficient of 0.190). Most enterocyte-specific antibodies (including sucrose isomaltase, SI) stain the apical surface, making colocalization studies challenging. However, given that enterocytes are the major cell type of the intestinal epithelium and that infected HIEs display an expanded and intense ORF2 signal (Fig. S6C), including cells that form the brush borders (as visualized by Phalloidin staining in Fig. 3E), it is most likely that enterocytes are targeted by HEV. Overall, the HEV ORF2 capsid protein was expressed in proliferating, absorptive as well as secretory cells.

### HEV-electroporated HIEs produce infectious HEV particles that are non-enveloped

We next infected HepG2/C3A cells with supernatant from HEV-electroporated HIEs (referred to as HIEs/HEV-3^G1634R^, collected on day 11 pe, Fig. 5A) in order to confirm the infectivity of the virions shed in the supernatants of HIEs/HEV-3^G1634R^. Indeed, the infected HepG2/C3A cells markedly expressed HEV ORF2 antigens on day 7 pi, resulting in a titer of 4.0×10^6^ focus forming units per mL (*N=3*). Moreover, viral replication was more efficient when proliferating 3D-HIEs were infected with HIEs/HEV-3^G1634R^ than with HepG2/C3A-HEV-3^G1634R^ (intracellular-derived virus stock, HEVenv^-^). In particular, at the peak of replication (day 2 pi), the viral load was 5.7 times higher when using the HIEs-derived HEV stock than when using the HepG2/C3A-derived HEV stock (*P=0.0679*) (Fig. 5B). To further characterize the type of HEV virions being released from the HIEs/HEV-3^G1634R^, HIEs electroporated with viral RNA were cultured in *in-house* organoid differentiation medium (bovine serum albumin-free) for 11 days, followed by the analysis of the ORF2 forms present in the supernatant by immunoprecipitation and western blot (Fig. 5C and D), and density gradient centrifugation (Fig. 5E). Using the P1H1 antibody that specifically recognizes the particle-associated ORF2i form^30^, an intense band corresponding to the ORF2i protein was detected in the HIEs/HEV-3^G1634R^ supernatant (Fig. 5C, left part in the top panel). Interestingly, the amount of ORF2i proteins in HIEs/HEV-3^G1634R^ supernatant was ∼1.4-fold higher than in the supernatant of human hepatoma PLC3 cells electroporated with HEV-3 RNA (‘PLC3/HEV-3’) control samples (Fig. 5C, right part in the top panel and 5D). Conversely, immunoprecipitation with the P3H2 antibody, which recognizes ORF2g/c forms^30^, resulted in poor detection of the secreted glycosylated ORF2g/c forms in the HIEs/HEV-3^G1634R^ supernatant, unlike in PLC3/HEV-3-derived virus (Fig. 5C, bottom panel and 5D). Density gradient analysis of the HIEs/HEV-3^G1634R^ supernatant showed high levels of HEV RNA in fractions 9, 10, and 11 that corresponded to HEV particles with densities of 1.15-1.19 g/cm^3^ (Fig. 5E), which are typical of non-enveloped particles^15,17^, thus indicating that viral particles released from HIEs/HEV-3^G1634R^ are mostly naked.

**Fig. 5.**
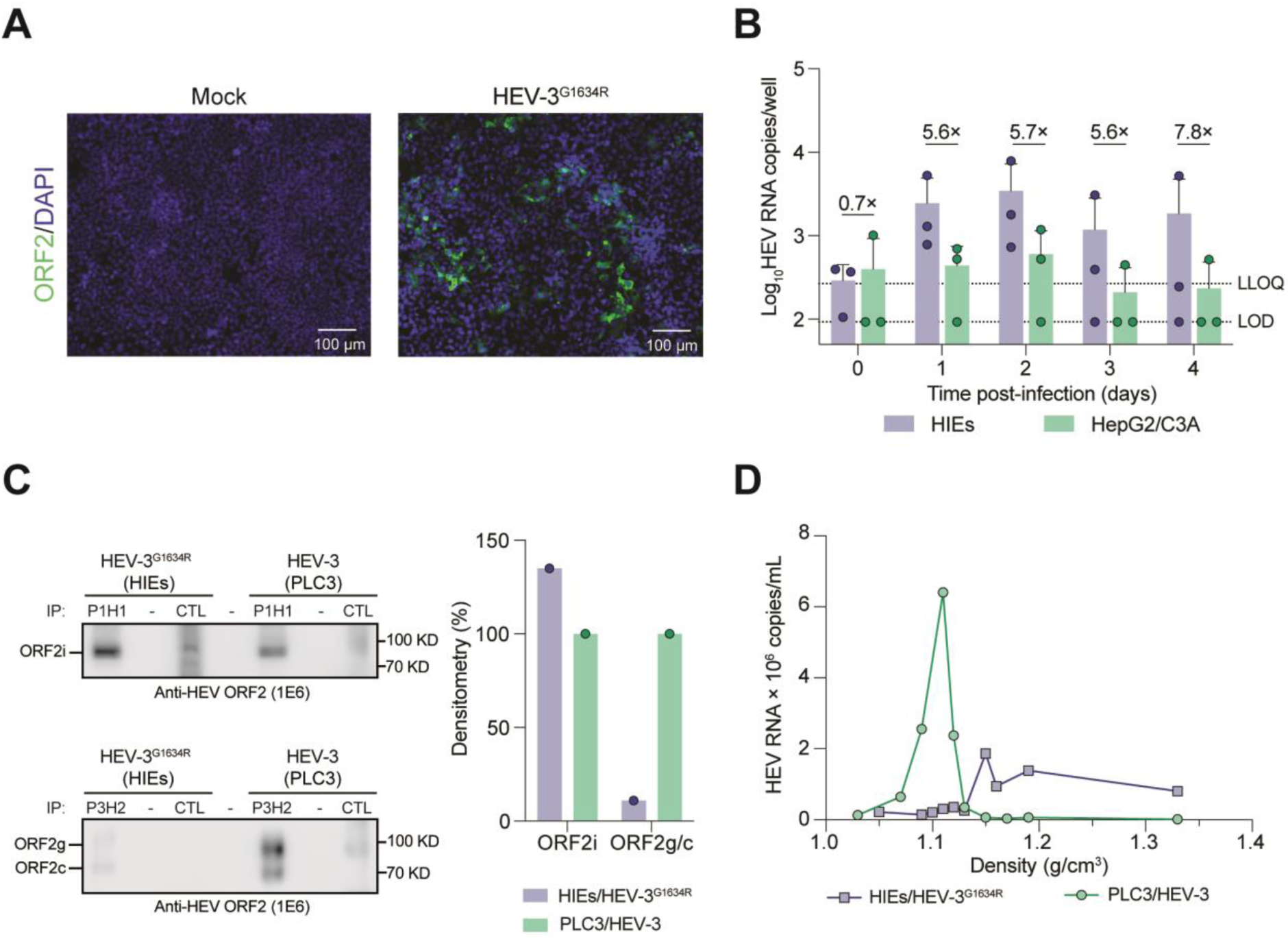
Characterization of HEV particles produced in HEV-electroporated HIEs. (A) Representative images of HEV ORF2 immunofluorescence staining in HepG2/C3A cells infected with the supernatant of HEV-3^G1634R^ electroporated HIEs (HIEs/HEV-3^G1634R^, 1:6 dilution) (*N=3*). Uninfected HepG2/C3A cells served as negative control. Scale bar – 100 μm. ORF2 (green); DAPI (blue). Images acquired on a DMi8 microscope (Leica) with a 10× objective. (B) Proliferating fetal 3D-HIEs were infected with 1.0×10^6^ GEs HEV produced in HIEs/HEV-3^G1634R^ or in HepG2/C3A cells (env^-^) (*N=3*). HEV RNA levels in the whole well were quantified by RT-qPCR. Day 0 pi represents 6h pi. Data are mean+SD. Fold changes in viral load were determined by comparing HIEs-derived infections against HepG2/C3A-derived infections at each time point (illustrated within the histogram). Statistical analysis was performed using the 2-way ANOVA, followed by Tukey‘s multiple comparisons test. (C) Expression of different forms of ORF2 protein (ORF2i, ORF2g, ORF2c) in supernatant of HIEs/HEV-3^G1634R^ and PLC3/HEV-3, as determined by immunoprecipitation using two different antibodies (P1H1 and P3H2) followed by western blot analysis. (D) Densitometry plot of band intensity, % HIEs compared to PLC3. (E) Density gradient of HIEs/HEV-3^G1634R^ and PLC3/HEV-3 supernatant. CTL, IP-negative controls.

## Discussion

Hepatitis E virus has been recognized as enterically transmitted since the late 1970s^31^, yet evidence supporting its infection of the gut has remained limited, which is partly due to its relatively inefficient *in vitro* cultivation. Understanding HEV infection in the primary infection site, the gut, as well as HEV dissemination to the liver is essential to grasp HEV pathogenesis. Additionally, identifying HEV’s intestinal cellular tropism should offer novel insights into virus-host interactions and potential disease mechanisms, which are highly relevant for an efficient clinical management and for the development of novel therapeutics. The development of human intestinal organoids has transformed the *in vitro* cultivation of various enteric viruses^32–34^, hence holding great potential for HEV, which is considered a slow-growing, hard-to-cultivate virus.

There are several methods to deliver infectious organisms into organoid cultures^35^. In this study, we explored three different ways to infect HIE cultures with HEV^29^. HEV was able to replicate in fetal and adult differentiated 3D-HIEs in an input-dependent manner. However, replication was somewhat limited, with a constant viral load detected until day 5 pi. Similar results were obtained when infecting 2D-differentiated HIE monolayers. The sustained high HEV viral load generated, despite the frequent medium changes, is comparable in terms of replication kinetics to what is observed in hepatocyte-like immortalized models^36^, primary hepatocytes^12^ and liver organoids^19^. Given that HEV is a slow replicating virus, prolonging the assay may likely result in a higher replication yield. However, this was not feasible because of the differentiated state of the cells and the consequent accumulation of debris in the lumen of the 3D organoid structure, resulting in loss of viability past this timepoint. In the 2D-HIE setup, HEV infection predominantly led to apical release of the virus, as demonstrated by a lower basolateral shedding. This indicates that access to the apical surface was not the limitation for efficient replication in 3D-HIEs. Moreover, these data, which are in line with previously described results using primary intestinal epithelial cells^17^, suggest that new HEV virions may be continuously excreted into the lumen.

Electroporation of single cell HIEs with HEV capped RNA proved to be the most effective method, leading to high viral yields, like with what is reported for human hepatocyte cell lines^12^. Hence, this approach is well suited for exploring the pathways that are involved in HEV infection of the gut. Many factors may be preponderant for successful HEV infection and/or further dissemination. One obvious advantage of electroporation is that it overcomes the early steps of entry and uncoating. Yet, a marked increase in viral load became evident only after three days pe. This observation suggests that the increase in viral load may be linked to slower processes such as cell proliferation or may be the result of multiple replication cycles after infection.

The model was able to confirm the enhanced replication of the HEV-3^G1634R^ variant, which contains a mutation in the RNA-dependent RNA polymerase, as it has been previously described in immortalized liver models^12^. We further validated the efficacy of the antiviral ribavirin in the HIE-HEV electroporation model. Given that the gut could serve as an important reservoir for HEV^17^, it is critical to evaluate (yet-to-be-developed) treatments against HEV in the gut. In this model, ribavirin effectively reduced viral RNA levels, both intra- and extracellularly, demonstrating that the model is amenable to evaluate the efficacy of novel therapeutic approaches in the intestinal compartment.

HEV-electroporated HIEs exhibited delayed growth (when compared to mock-infected), but an increase in the number of proliferative cells was observed. While mature cell types are present in all HIE culture types (including 3D- and 2D-HIEs), only the undifferentiated HIE single-cells had an elevated proportion of proliferative cells (*i.e*. approximately 40%). Moreover, the fact that nearly 70% of HEV-infected cells were proliferative cells indicates that this cell type better supports HEV replication, when compared to mature cell types. Of note, the fact that electroporation of single-cell HIEs is performed is an important factor that facilitates the first round of infection. However, the increase in viral load over 11 days, in parallel with increasing numbers of proliferative cells during infection, is highly suggestive that infection of this cell type is a determining factor for HEV sustained and highly efficient replication in these cultures. It would be relevant to directly compare this to electroporation of single-cell differentiated HIEs but these cells are, however, not viable after electroporation.

The infection of mature cell types may have additional implications regarding HEV infection and disease. Infection of enterocytes is a finding that is not surprising as HEV can replicate in Caco-2 cells^37^. Enteroendocrine cells are estimated to comprise approximately 1% of the intestinal epithelial cell population, serving as sensory sentinels of the intestinal environment and possessing rich endocrine functions^38^. HEV infection of enteroendocrine cells could explain the early symptoms in HEV-infected individuals, such as vomiting^14^. Similar findings were reported for the enteric human rotavirus, which causes acute gastroenteritis and also infects enteroendocrine cells, inducing serotonin release to the basolateral side of the epithelium, which consequently stimulates vagal afferent nerves and the vomiting center in the brain^39,40^.

One important aspect in HEV biology is the characterization of the shed virions from infected tissues to determine whether they are naked or quasi-enveloped. Density gradient centrifugation showed that infectious HEV virions released by infected HIEs were naked. Since naked virions are associated with a higher infectivity^12^, their presence likely facilitated further dissemination of progeny virions to new cells, hence explaining the higher viral titer of the HEV stock derived from electroporated HIEs as well as the higher amounts of expressed ORF2i proteins. Moreover, infection of 3D-HIEs with HIEs/HEV-3^G1634R^ yielded a higher virus replication, most likely due to the higher infectivity of the inoculum that mainly contains naked virions. This finding is novel and contrasts with a previous study that detected mainly quasi-enveloped virus shed by human primary intestinal cells^17^. Likewise, HEV egress from liver cell cultures is reported to occur in the quasi-enveloped form^41^.

We here postulate that while HEV can infect virtually all intestinal cell types, infection of proliferative cells is key for a more productive and sustained infection of the gut, likely due to an efficient production of naked virions in these cells. This is based on the following observations: (i) only HIEs cultures used for electroporation present high proportion of proliferative cells; (ii) a marked increase in HEV RNA levels after electroporation happened over a period of 11 days, implying that a more efficient multiple-replication cycle process is taking place; (iii) the virions produced in electroporated HIEs are mainly naked, which supports the notion of a more efficient replication, also because re-infection of new HIEs happens with rapid replication kinetics (*i.e.* peaking at 48 h and yielding higher HEV RNA levels). In contrast, when differentiated HIEs in 3D or 2D were used (where few to no proliferative cells are left), lower and constant HEV RNA levels were reached. Likewise when human intestinal primary cells were used by Marion *et al.*^17^, cultures likely did not contain proliferative cells anymore, thereby yielding quasi-enveloped virions to be produced. Therefore, efficient production of naked virions (which can better infect new cells) is a key feature to obtain an overall increase in viral load overtime; a process not occurring in cultures in which insufficient proliferative cells are present and infected.

This hypothesis also suggests that virus progeny produced by HEV-infected mature (non-proliferative) cell types may consist mainly of quasi-enveloped virions, which are less efficient in infecting new cells and thus resulting in lower and steady HEV RNA levels over time. Further studies are needed to understand how intracellular processes differ between infected proliferative and non-proliferative intestinal cells. Along the same line, additional studies utilizing different intestinal segments from various donors would provide a more thorough characterization of HEV replication and shedding in the intestine.

In conclusion, we established an efficient infection model for HEV using intestinal organoids. Direct delivery of HEV RNA to single-cell HIEs that continued to develop into 3D-HIEs resulted in high levels of infectious naked virions. We discovered that HEV is able to infect multiple types of intestinal cells, mainly proliferative cells. By contrast, lower HEV RNA yields were detected when fewer or no proliferative cells were present and infected. Thus, the fast epithelial cell turnover of the gut, with direct infection of proliferative cells, is key for establishing a highly productive HEV infection of the gut, likely impacting viral dissemination within and beyond this organ towards the liver. Overall, this work demonstrates the relevance of the gut as a potential HEV reservoir, suggesting that part of the naked HEV shed in the feces originates directly from this organ.

## Material and Methods

### Human intestinal enteroid culture

Human intestinal enteroids (HIEs) derived from fetal ileum (HT124), adult jejunum (J2) or adult small intestine (Lonza), were maintained at 37°C with 5% CO_2_ in extracellular matrix (Matrigel, Corning) and IntestiCult™ Organoid Growth Medium (OGM, StemCell Technologies), that was replaced every other day. Differentiation was triggered by Wnt3a removal with IntestiCult™ Organoid Differentiation Medium (ODM, StemCell Technologies) for 5 days with medium changes every other day. HIEs culture and experimentation was performed under the approval of the Ethical Committee comite KU Leuven (approval number G-2024-8519-R2(AMD).

### Viruses

HEV-3 full-length Kernow-C1 p6 infectious cDNA clone (GenBank accession number JQ679013) and its G1634R variant, containing the G1634R mutation in the viral polymerase, were previously detailed^6^. The subgenomic Kernow-C1 p6/luc, with part of ORF2 replaced by the *Gaussia* luciferase gene, was constructed as described previously^42^. Similarly, the genotype 1 reporter replicon Sar55/S17/luc was derived from the HEV strain Sar55/S17 (GenBank accession number AF444002), as described earlier^43^. The construction of plasmid pLA-B350/luc was previously reported^44^.

### Enteroid electroporation

HIE culture and preparation of single cell suspension was performed as described previously^45^. Briefly, 3D-HIEs were kept in OGM for five days. Forty-eight hours prior to electroporation, medium was replaced by ODM supplemented with 5 µM CHIR99021 (StemCell technologies) and 10 µM Y-27632. Twenty-four hours prior to electroporation, medium was replaced by ODM supplemented with 5 µM CHIR99021, 10 µM Y-27632 and 1.25% (V/V) DMSO. HIE single cell suspensions were prepared using TrypLE express supplemented with 10 µM Y-27632 for 20 min at 37°C. Single-cell suspensions of HIEs (1.0 × 10^6^ cells) were resuspended in 200 μL BTX buffer (BTXpress; BTX Harvard Apparatus), mixed with 10 μg capped Kernow-C1 p6, Kernow-C1 p6-G1634R, Kernow-C1 p6/luc, Kernow-C1 p6 G1634R/luc, pLA-B350/luc, or Sar55/S17/luc RNA in a 4-mm (VWR International) cuvette. An ECM 830 Electro Square Porator™ (BTX Harvard Apparatus) was used to deliver two pulses at 450 V for 2 msec with 100 ms interval between each pulse. The addition of viral RNA to the electroporation mixture was omitted for the cell control samples. Electroporated cells were incubated in ODM with 10 µM Y-27632 for 40 min at RT, then centrifuged and resuspended in 100% matrigel and plated in pre-warmed 48-well plates (20 µL drop containing 1.0 × 10^5^ cells). For immunofluorescence experiments, electroporated cells were resuspended in 50% matrigel-ODM solution and plated in black wall 96-well plates (Greiner) pre-coated with 5% matrigel. Electroporated cells were cultured in ODM supplemented with 10 µM Y-27632, 5 µM CHIR99021 (StemCell technologies) and 1.25% V/V DMSO), either with or without ribavirin (Sigma-Aldrich).

**Additional methods are provided in the Supplementary information**.

## Abbreviations

CHGA: Chromogranin A
HEV: hepatitis E virus
HIEs: human intestinal enteroids
HIOs: human intestinal organoids
Lgr5+: leucine-rich-repeat-containing G-protein-coupled receptor 5
ODM: organoid differentiation medium
OGM: organoid growth medium
ORF2: open reading frame 2
pe: post-electroporation
pi: post-infection
3D: three-dimensional
2D: two-dimensional
WGA: wheat germ agglutinin

**GenBank accession number** JQ679013, AF444002, KM516906.

## Financial support

This work was partly supported by the Horizon 2020-funded ITN program OrganoVIR (Grant 812673) to N.S.F.; J.R.P. was supported by the KU Leuven Starting Grant STG/21/028. X.Z. was supported by the China Scholarship Council (grant no. 201906170033). L.Cor. was supported by the ANRS-MIE (ECTZ246976). J.V.D. was supported by the EU (DT-NMBP-23). W.C. research was performed using the “Caps-It” research infrastructure (project ZW13-02) that was financially supported by the Hercules Foundation (FWO) and Rega Foundation, KU Leuven.

## Acknowledgements

We thank Prof. Luc Verschaeve (formerly at the Scientific Institute of Public Health, Belgium), Prof. Jason Spence (University of Michigan Medical School, United States) and Prof. Mary Estes (University of Michigan Medical School, United States) for providing the HepG2/C3A cells, fetal ileum HIEs and adult jejunum HIEs, respectively. We thank Prof. Suzanne Emerson (formerly at the National Institute of Allergy and Infectious Diseases, National Institutes of Health, United States) for kindly provide the plasmids Kernow-C1 p6, Kernow-C1 p6/luc, and Sar55/S17/luc. We also thank Prof. Eike Steinmann (Ruhr University, Germany) for the anti-ORF2 HEV-specific rabbit hyperimmune serum. We thank Joost Schepers (KU Leuven, Belgium) for the kind assistance with the high-content imaging analysis. We truly thank Dr. Martin Ferrié (KU Leuven, Belgium) for all the insightful discussions.

## Conflict of interest statement

No potential competing interests were disclosed by the authors.

## Data Transparency Statement

The appropriate datasets will be made publicly available upon publication.

## Notes

### Competing Interest Statement

The authors have declared no competing interest.

### Summary of Updates

New data as been added to support the hypothesis that the proliferative cell niche plays an important role in HEV infection of the gut. Moreover, important finding have been included such as the characterization of the HEV viral particles produced in HIEs.

