## Supplementary information for "Proliferative cell targeting and epithelial cell turnover fuels hepatitis E virus replication in human intestinal enteroids"

Table of contents

Supplementary Materials and Methods ------------------------------------------------------- 2

Supplementary Figures ---------------------------------------------------------------------------- 9

Supplementary References -------------------------------------------------------------------- 17

### Material & methods

*Cells*

HepG2/C3A cells (ATCC CRL-10741) were cultivated in Dulbecco's Modified Eagle Medium (DMEM, Gibco), supplemented with 10% inactivated fetal bovine serum (FBS, Gibco), 2 mM L-glutamine (Gibco), 0.75 g/L sodium bicarbonate (Gibco) and 1 mM sodium pyruvate (Gibco). For seeding of HepG2/C3A cells, plates were coated with 100 µg/mL rat tail collagen (Sigma-Aldrich) and 0.02 M glacial acetic acid in PBS for a minimum of 2h at 37°C. Subsequently, the plates were washed three times with PBS. PLC3, a subclone of PLC/PRF/5 hepatoma cells (CRL-8024), was cultured as previously described[1].

*Generation of infectious virus stocks*

Capped viral RNA was generated from the cDNA plasmids as previously described[2]. In brief, Kernow-C1 p6, Kernow-C1 p6 G1634R, Kernow-C1 p6/luc, and pLA-B350/luc were linearized by MluI (Promega), while Sar55/S17/luc was linearized by BglII (Promega). Subsequently, *in vitro* transcription was carried out using the Ribomax large-scale RNA production systems (Promega), and subsequent capping was performed using the ScriptCap m7G capping system (Cellscript), according to the manufacturer's protocols.

Kernow-C1 p6 or G1634R virus stocks were either a mixture of non-enveloped and enveloped virus ($env^{+/-}$), or only non-enveloped virus ($env^{-}$)[3,4]. In short, capped Kernow-C1 p6 or G1634R RNA was transfected into HepG2/C3A cells through electroporation as previously described[3,4]. After seven days of incubation, extracellular enveloped and intracellular non-enveloped Kernow-C1 p6 or G1634R were harvested as follows. For extracellular virus, supernatant was collected and centrifuged at 4,000 rpm at 4°C for 15 min, followed by the removal of cell debris, the concentration of the virus in the supernatant using minimate tangential flow filtration (100K pore size capsules, Pall Corporation), according to the manufacturer's protocol. For intracellular virus, electroporated cells were collected by trypsinization, resuspended in fresh complete DMEM, and subjected to three freeze-thaw cycles in liquid nitrogen to release the virus from the cells. The lysate was subsequently centrifuged at 10,000 *xg* for 10 min. Non-enveloped virus either or not mixed with concentrated enveloped virus was aliquoted and stored at -80°C until further use. Levels of infectious virus and viral RNA in the stock were determined by immunofluorescence staining using the HEV ORF2 protein and RT-qPCR, respectively, as previously described[4]. Extracellular Kernow-C1 p6 virus from electroporated PLC3 cells was generated as previously described17 and used for the immunoprecipitation and gradients.

*HEV infection in 3D enteroids*

HEV infection of 3D-HIEs was performed as previously described for other enteric viruses with minor modifications[5–7]. Briefly, 3D-HIEs were removed from matrigel, thoroughly washed with CMGF(-) medium (Advanced DMEM/F12 (Gibco) supplemented with GlutaMAX-1 2 mM (Gibco), HEPES 10 mM (Gibco), penicilin-streptomicin 100 U/mL (Gibco) and primocin 100 µg/mL (Invivogen), and gently fragmented. 3D-HIEs were resuspended in infection medium [ODM supplemented with 500 µM glycochenodeoxycholic acid (GCDCA; Sigma-Aldrich) and 10 µM Y-27632 (StemCell Technologies)] and inoculated with HEV or mock-infected. After virus inoculation, 3D-HIEs were incubated for 6h at 37°C with homogenization every hour. After, 3D-HIEs were washed with CMGF(-) (5x), resuspended in matrigel and plated in pre-warmed 48-well plates (20 µL drop containing $1.0 \times 10^5$ cells). After matrigel polymerization for 10 min at 37°C, infection media [ODM supplemented with 500 µM GCDCA, 10 µM Y-27632 and 2 mM ruxolitinib (Toronto Research Chemicals)] was added to each well. At multiple timepoints post infection (pi), cells and supernatant were collected for RNA extraction. Infection of undifferentiated 3D-HIEs followed the same protocol with HIEs always cultured in OGM, including the infection medium.

*HEV infection in transwell*

Proliferating 3D-HIEs were collected and dispersed into single cells using TrypLE express (Gibco) and seeded on semipermeable transwell inserts (0.4 µm PET membrane, VWR) that were precoated with collagen type IV (EMD Millipore). After 2 days in OGM, cells were maintained in ODM until formation of a polarized monolayer (6-7 days). TEER of monolayers was measured using an epithelial voltohmmeter (EVOM, WPI). Polarized monolayers were infected with HEV Kernow-p6-G1634R through the apical side for 6h at 37°C. After, cells were washed five times with CMGF(-). ODM supplemented with 500 µM GCDCA and 2 µM ruxolitinib was added to the apical and basal compartments and incubated at 37°C and 5% $CO_2$. Supernatant from the apical and basolateral sides, and cells from the apical side were harvested at multiple time points pi. Infection of undifferentiated 2D-HIE monolayers followed the same protocol with HIEs always cultured in OGM, including the infection medium.

*Luminescence-based replicon assay*

To track the replication kinetics of luciferase expression activities in HIEs electroporated with HEV replicons carrying the luciferase gene, 20 µL of culture medium was collected at various time points post-electroporation (pe). Luciferase activity was assessed via the Renilla luciferase kit (Promega). In brief, 20 µL of the culture medium was transferred to white 96-well Culture Plates (PerkinElmer), followed by the addition of 50 µL (1:100 diluted) *Gaussia* substrate. Luminescence produced by the secreted *Gaussia* luciferase was measured using a Spark microplate reader (Tecan).

*Cell viability*

Enteroid viability was assessed using the Live-Dead Cell Viability Assay Kit (EMD Millipore) according to the manufacturer's instructions. Briefly, media was aspirated and replaced with dye mixture and incubate for 60 min at 37°C. Images were acquired on a DMi8 (Leica) fluorescence microscope with a 10x objective.

*RNA extraction and quantitative real-time PCR (RT-qPCR)*

Extracellular viral RNA was extracted from 50 or 150 µL supernatant collected from infected/transfected HIEs or HepG2/C3A cells, by using the NucleoSpin RNA virus kit (Macherey-Nagel). For intracellular viral RNA, cells were harvested in TRI Reagent and RNA extracted with DirectZol RNA extraction kit (Zymo Research, R2051) according to manufacturer's instructions. HEV RNA loads were quantified using an established human HEV RT-qPCR protocol as previously described[8,9].

HEV RNAs for HEV particle characterization were extracted from 140 µL culture supernatants using the QiAmp viral RNA mini kit (Qiagen). HEV RNA levels were quantified by RT-qPCR using the Takyon One-Step qPCR kit and using primers (5'-AAGACATTCTGCGCTTTGTT-3' (F) and 5'-TGACTCCTCATAAGCATCGC-3' (R)) and a probe (5'-FAM-CCGTGGTTCCGTGCCATTGA-TAMRA-3') targeting a conserved region of ORF1. RT-qPCR was performed on a QuantStudio 3 device.

Gene expression analysis of intestinal epithelial cell type mRNA levels was performed in samples harvested in TRI Reagent and extracted with DirectZol RNA extraction kit. An iTaq Universal SYBR Green One-Step Kit (Bio-Rad) was used to quantify the relative expression of different intestinal cells markers: Lgr5 (stem-cells), Lysozyme (Paneth cells), sucrose isomaltase (mature enterocytes), mucin 2 (goblet cells) and chromogranin A (enteroendocrine cells), using specific primers as previously described[5]. RT-qPCR was carried out in a QuantStudio 5 Real-Time PCR System (Applied Biosystems). GAPDH was used to normalize gene expression. Relative expression was determined using the $\Delta\Delta$Cq method.

*Immunofluorescence staining and focus-forming assay*

Immunofluorescence assays were performed at the peak of replication, *i.e.,* after day 11 pe, as described previously[7]. Briefly, supernatant was removed without disturbing the HIE layer and cells were fixated with a 2% paraformaldehyde (PFA) solution at 4°C overnight. Then, cells were permeabilized (0.2% Triton X-100), blocked (1% goat serum and 3% bovine serum albumin (BSA) and incubated overnight with primary antibodies at 4°C. Subsequently, cells were incubated with corresponding secondary antibodies (Invitrogen), followed by nuclear counterstaining with DAPI (4',6 Diamidino 2 Phenylindole, Dilactate, Invitrogen) for 1h at RT. The following antibodies or dyes were used: anti-ORF2 (HEV-specific rabbit hyperimmune serum), anti-Ki67 (EMD Millipore, MAB4190), anti-sucrose isomaltase (Santa Cruz Biotechonology, sc-393424), anti-chromogranin A (Santa Cruz Biotechonology, sc-393941), phalloidin (Invitrogen, A12380) or wheat germ agglutinin (WGA, Vector laboratories, RL-1022). Images were acquired on an Andor Dragonfly 200 series High Speed confocal platform system (Oxford instruments, Abingdon, UK) at a 25× magnification connected to a Leica DMi8 microscope (Leica Microsystems, Wetzlar, Germany). Image processing was done with the Imaris analysis software (version 9.8.2). The 3D colocalization analysis of virus and intestinal epithelial cell markers was done using Imaris 3D colocalization software (voxel-based colocalization). Images are presented as maximum projections.

For high-content imaging (HCI) experiments, immunofluorescence staining was performed using DAPI (blue), HEV ORF2 (green), Ki67 (red), CHGA (red) and Phalloidin CF660R (far-red). Confocal microscopic images of the HIEs were acquired using an Operetta CLS High-Content Imaging and Analysis system (Revvity). A 20X water immersion objective was used to capture 16 planes in four different channels with an overal z-height of 15 µm. Image analysis was performed on the maximum projection of the z-stacks using the Harmony® software (Revvity). The phalloidin stained cell membrane was used to identify the HIEs and downstream image analysis algorithms were used for the individual HIE segmentation and morphological feature extraction. Cell count was performed by counting the DAPI positive nuclei. Proliferative and enteroendocrine cells were counted based on positive Ki67 and CHGA signals, respectively, in close proximity of the DAPI stained nuclei. The presence of viral particles was detected by positively stained ORF2 and infected objects (HIEs, proliferative or enteroendocrine cells) were identified when both channels were positive.

For detecting HEV ORF2 in HepG2/C3A cells infected with HEV stocks, HepG2/C3A cells (7500 cells/well) were seeded in a 96-well plate with 10% complete DMEM the day prior to HEV infection. Viruses were serially diluted at 1:3, with a 1:6 dilution as the starting dilution. On day 7 pi, the cells were fixed with 3.7% PFA and incubated for 30 min at RT. Next, cells were permeabilized (0.1% Triton X-100) and blocked (3% BSA). Subsequently, the cells were incubated overnight at 4°C with primary antibody anti-ORF2 (HEV-specific rabbit hyperimmune serum), followed by incubation with the secondary antibody goat anti-rabbit Alexa Fluor 488 (1:800, Thermo Scientific) and nuclear counterstaining with DAPI (1:800; Sigma) for 1h. Fluorescent images of each well were acquired using a high-content imager (Arrayscan XTI, Thermo Fisher Scientific). Viral titers in focus-forming units were determined by counting the number of single HEV ORF2-positive cells through image analysis using the HCS Studio Cellomics software (Thermo Fisher Scientific, Version 6.6.2).

*Immunoprecipitation (IP)*

Antibodies recognizing the different forms of the HEV ORF2 protein, either P1H1 antibody recognizing the ORF2i form or P3H2 recognizing the three ORF2 forms (ORF2i, ORF2g, ORF2c) were used[10]. These antibodies as well as an IgG control antibody, were coupled to Epoxy M-270 Dynabeads beads (Thermofisher) overnight at 37°C. Beads were washed and incubated with either 0.5%-Triton treated supernatant (IP-P1H1 and an appropriate IgG control (Santa cruz biotechnology, sc-2025) or heat-inactivated supernatant (IP-P3H2 and an appropriate IgG control) for one hour at RT. Beads were washed six times and heated in Laemmli buffer. Proteins were separated by 10% SDS-PAGE and transferred onto nitrocellulose membranes, where detection of ORF2 proteins was done by using 1E6 monoclonal antibody (Millipore) and corresponding peroxidase-conjugated secondary antibodies, followed by exposure using the ImageQuant 800 chemiluminescent imaging system (Cytiva Life Sciences).

*Density gradient analysis*

Density gradient analysis was performed as previously described[1]. In brief, supernatants with equal HEV RNA copy numbers from PLC3/ Kernow-C1 p6 and HIEs/ Kernow-C1 p6 G1634R were layered on a preformed 7.5% to 40% iodixanol gradient, followed by centrifugation for 16 hours at 160,000×*g* at 4°C, using a SW41 swinging-bucket rotor (Beckman Coulter). Twelve fractions of 1 mL were collected, and their density was measured by refractometry. The HEV RNA load was quantified by RT-qPCR as described above.

*Statistical analysis*

Statistical analysis was performed using GraphPad Prism 10.2.0 (GraphPad Software, Inc.). All data were presented as mean + SD of at least 3 independent experiments unless otherwise stated. The specific methods of statistical analysis and *P* values are indicated in the figure legends or the text.

### Supplementary Figures

**Supplementary Fig. 1**

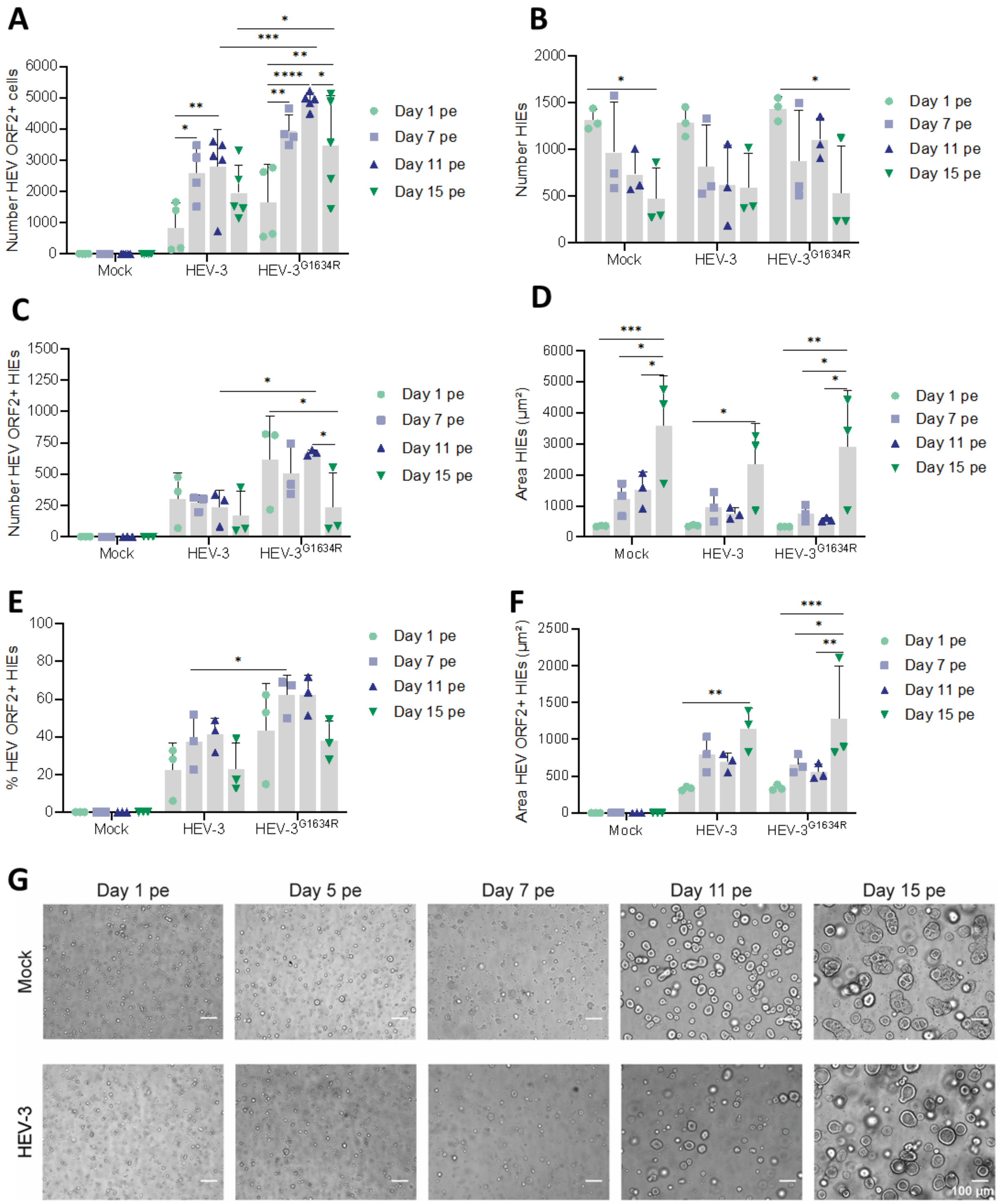


**Fig. S1. Effect of HEV infection in HIE growth.** (A) Number of HEV ORF2 positive cells, (B) number of HIEs, (C) number of HEV ORF2 positive HIEs, (D) HIEs size (µm²), (E) percentage of HEV ORF2 positive HIEs and (F) size of HEV ORF2 positive HIEs ($\mu m^2$) in mock-, HEV-3 or HEV-3$^{G1634R}$- electroporated HIEs at day 1, 7, 11 and 15 pe. Number of cells and HEV ORF2 positive cells were determined by high-content imaging of cell nuclei (DAPI) and HEV ORF2. Number, percentage and area of HEV ORF2 positive HIEs were determined based on actin (phalloidin) and HEV ORF2 staining and identified using HCI. Clustered HIEs were segmented using in-house image analysis algorithms prior to size determination. Statistical analysis was performed using the 2-way ANOVA, followed by Tukey's multiple comparisons test. *, $P \leq 0.05$; **, $P \leq 0.01$; ***, $P \leq 0.001$; ****, $P \leq 0.0001$. (G) Representative bright-field images of mock and HEV-3 electroporated fetal HIEs at day 1, 5, 7, 11 and 15 pe. Images acquired on a DMi8 microscope (Leica) using a 10× objective. Scale bar - 100 µm. pe, post-electroporation.

**Supplementary Fig. 2**

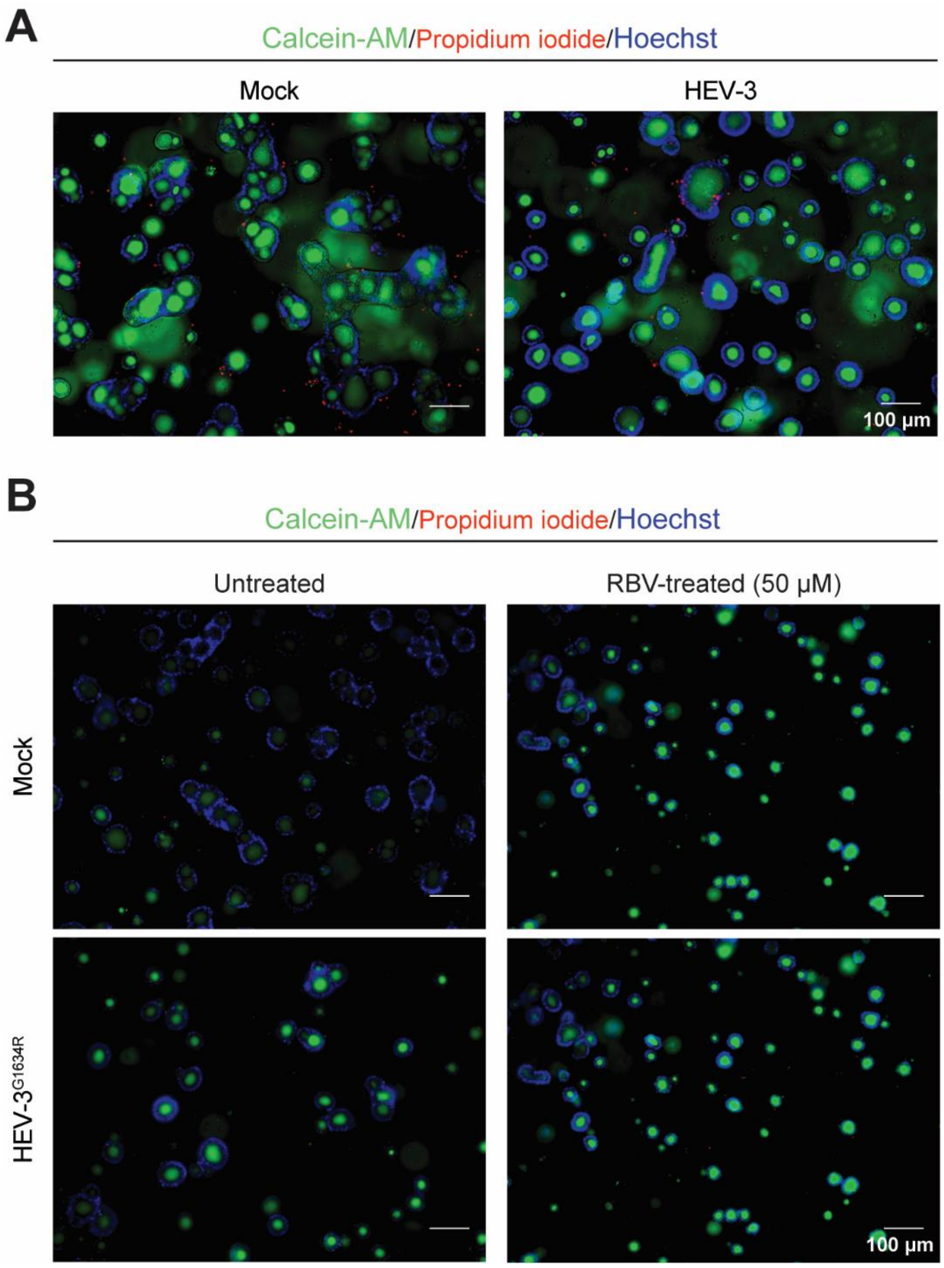


**Fig. S2. Viability of HEV electroporated fetal HIEs**. (A) Representative images of mock and HEV-3 electroporated fetal HIEs at day 15 pe. (B) Representative images of mock and HEV-$3^{G1634R}$ electroporated fetal HIEs at day 15 pe, with or without RBV (50 µM) treatment. Green - Calcein-AM. Red - propidium iodide. Blue - Hoechst 33342. Images acquired on a DMi8 microscope (Leica) with a 10× objective. Scale bar – 100 µm. pe, post-electroporation.

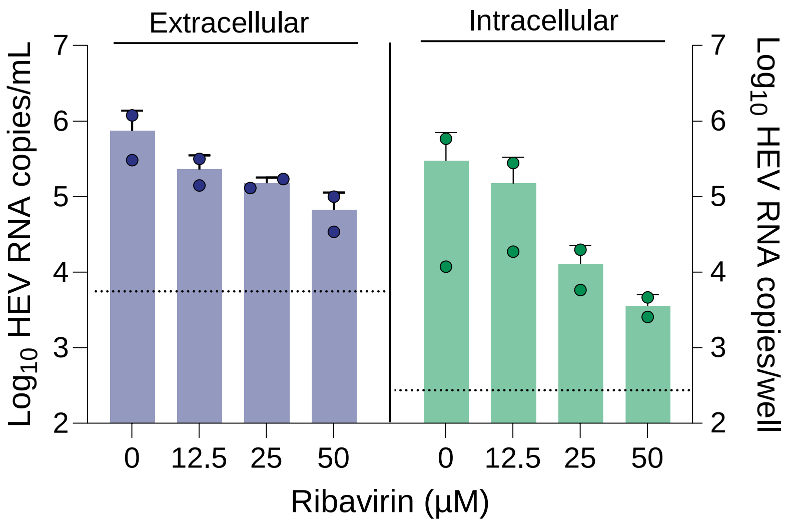


**Fig. S3. Ribavirin blocks HEV replication in HIEs.** HEV-3$^{G1634R}$ electroporated fetal HIEs treated with different concentrations of RBV (0, 12.5, 25 or 50 µM) (*N=2,* performed in duplicate). Viral load was determined at day 13 pe by RT-qPCR in HIE cell lysates or in 50 µL supernatant. Dotted line represents the lowest limit of quantification (LLOQ). pe, post-electroporation; RLU, relative luminescence unit; RBV, ribavirin;. Data are mean + SD.

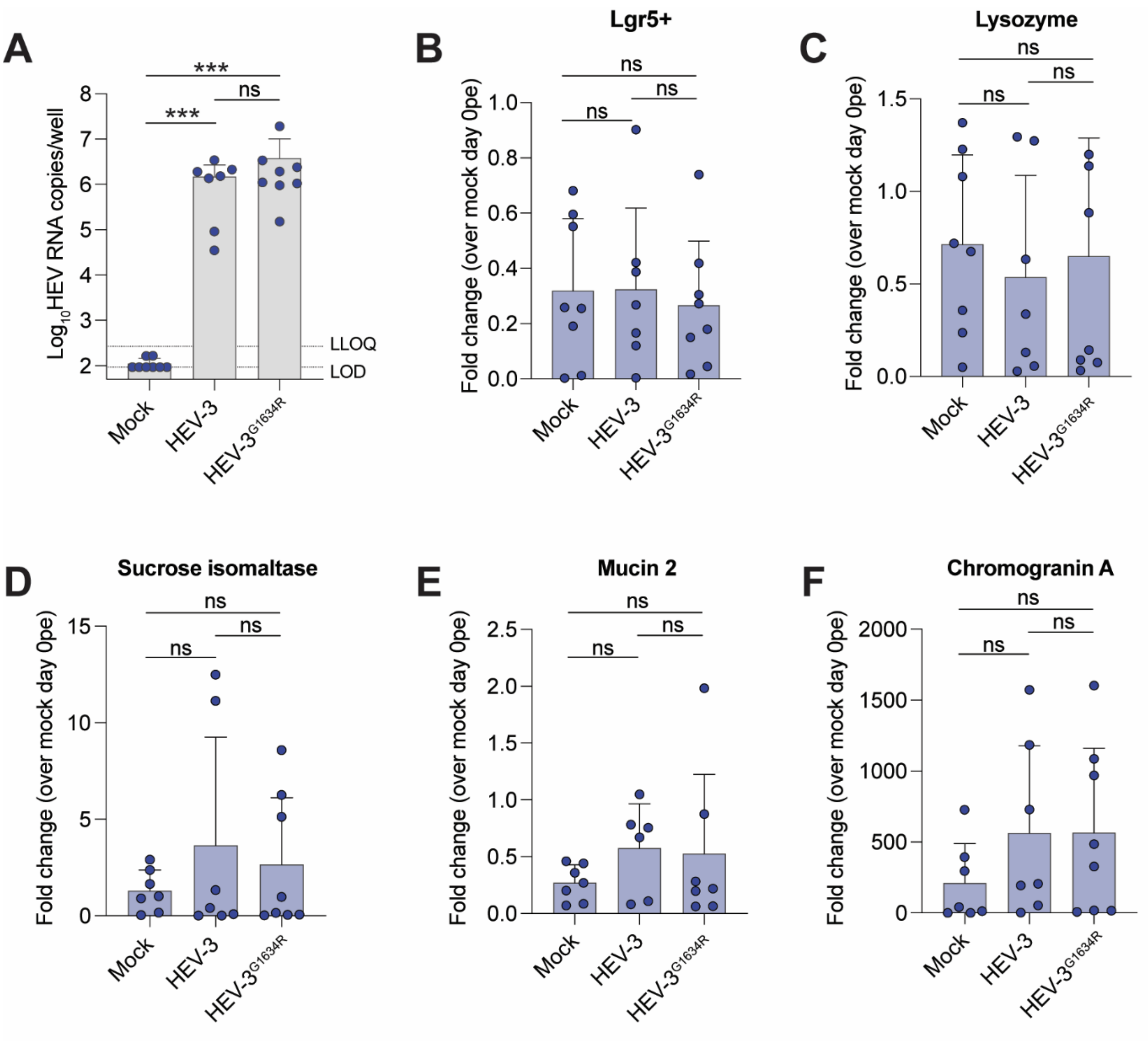


**Fig. S4. Intestinal epithelial cell expression upon HEV electroporation of HIEs.** (A) HEV viral RNA levels in HIEs electroporated with mock, HEV-3 or HEV-3$^{G1634R}$ capped RNA on day 11 pe. HEV RNA of the whole well (pool of supernatant and cell lysate) was quantified by RT-qPCR. (B to F) Gene expression of stem cells (Lgr5$^{+}$), Paneth cells (lysozyme), enterocytes (sucrose isomaltase), goblet cells (MUC2) and enteroendocrine cells (chromogranin A) in fetal HIEs electroporated with mock (*N*=8), HEV-3 (*N*=7) or HEV-3$^{G1634R}$ (*N*=8) capped RNA at day 11 pe, as quantified by RT-qPCR. Cell number was normalized based on GADPH gene expression. Relative expression was determined using the ΔΔCq method. Fold change is relative to mock HIEs at day 0 pe. Statistical analysis was performed using Grubbs test to remove significant outliers (*P* <0.05) and Mann Whitney test. ***, *P* ≤0.001; ns, not significant.

**Supplementary Fig. 5**

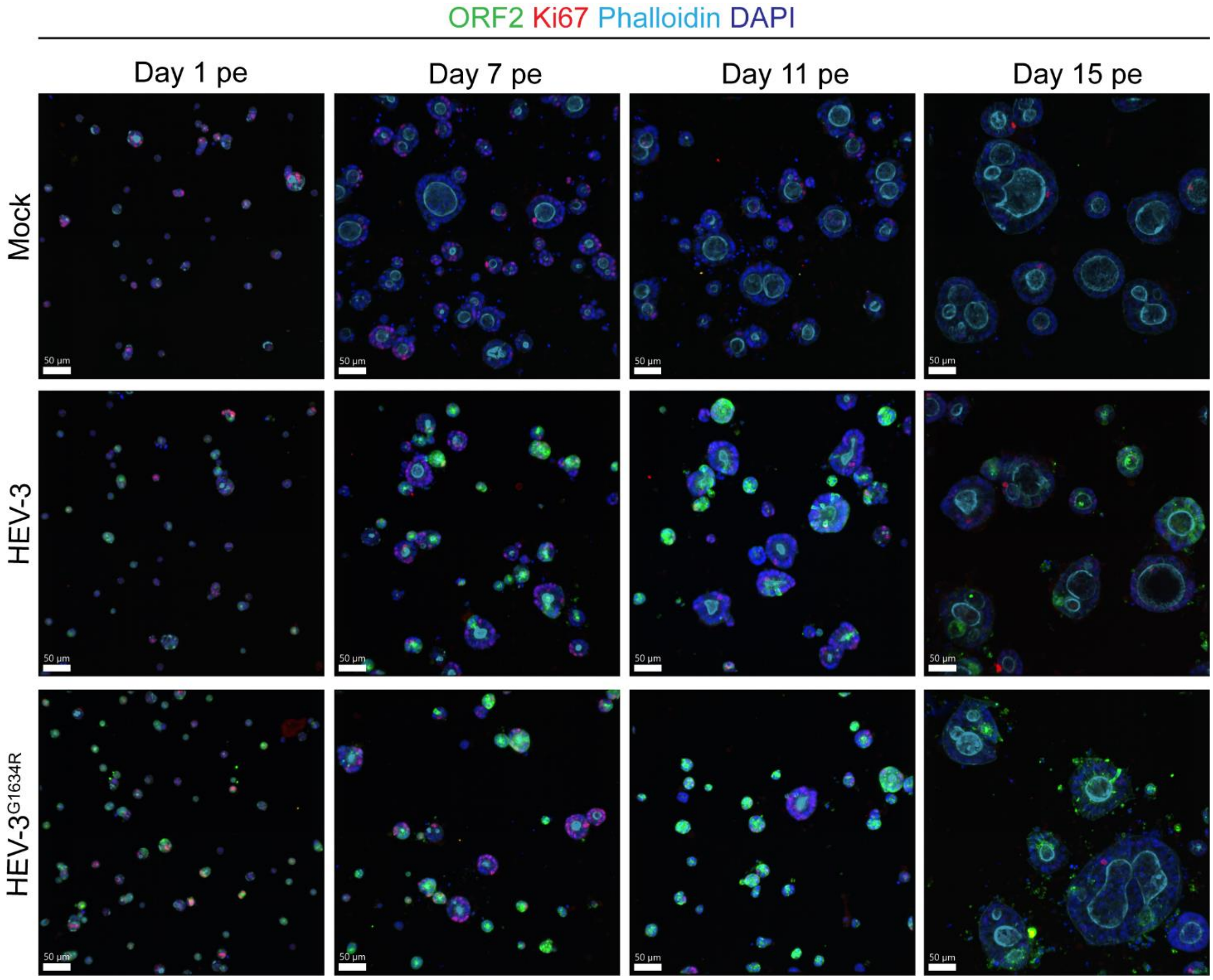


**Fig. S5. HEV infection of proliferating cells.** Representative images of immunofluorescence staining of mock, HEV-3 or HEV-3$^{G1634R}$ electroporated fetal HIEs at day 1, 7, 11 and 15 pe. Cells were stained with HEV ORF2-specific antibody (green), proliferating marker Ki67 (red) and actin (phalloidin, cyan). DAPI (blue) was used as counterstaining (nuclei). Scale bar – 50 μm. pe, post-electroporation.

### Supplementary Fig. 6

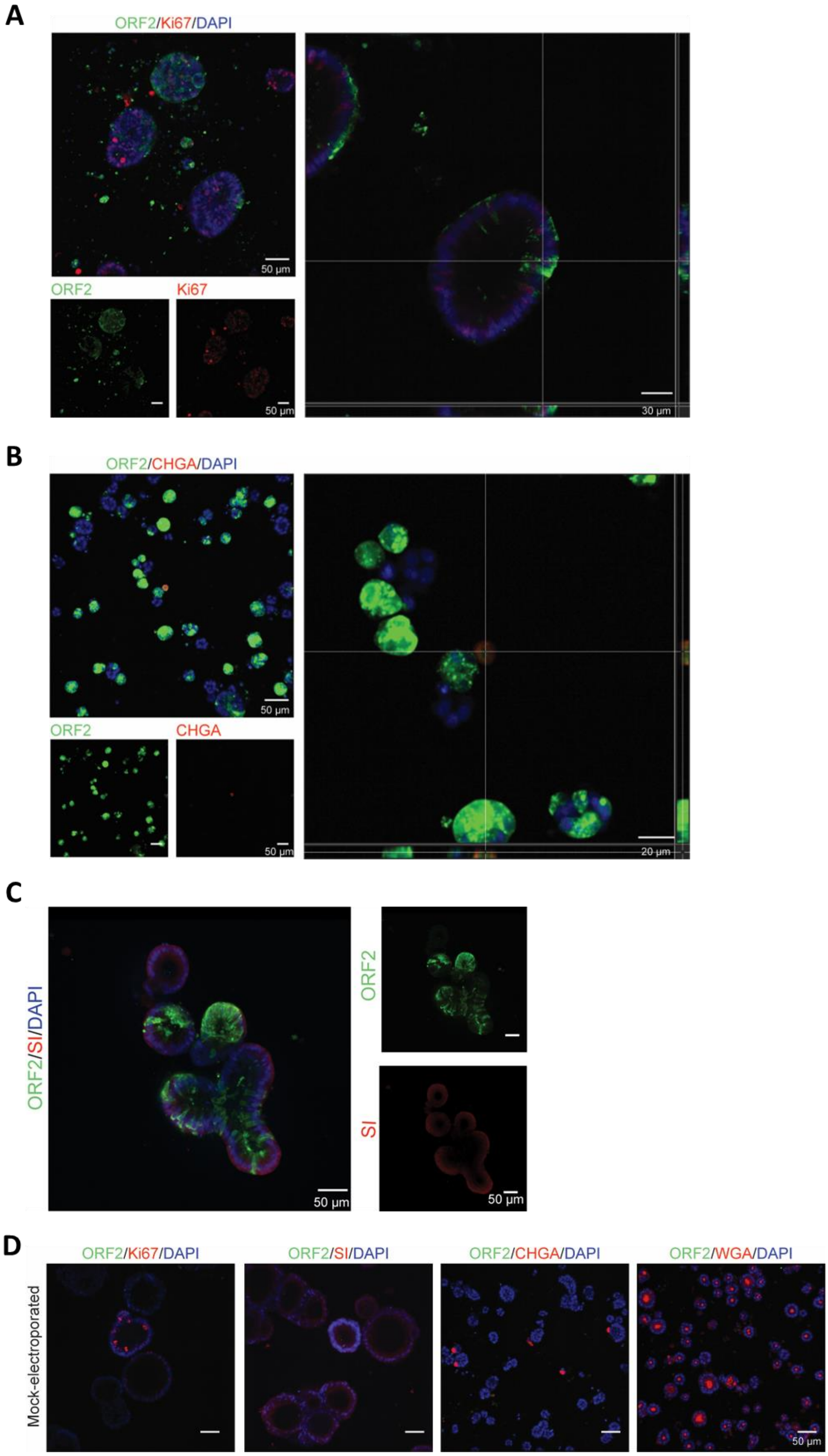

**Fig. S6. Tropism of HEV in HIEs.** (A) Presence of HEV capsid protein in proliferating cells. Immunofluorescence staining of HEV ORF2 (green) and proliferating cells (Ki67, red) in HEV-$3^{G1634R}$electroporated HIEs at day 11 pe. Counterstaining with DAPI. (B) Presence of HEV capsid protein in enteroendocrine cells. Immunofluorescence staining of HEV ORF2 (green) and enteroendocrine cells (CHGA - chromogranin A, red) in HEV-$3^{G1634R}$ electroporated HIEs at day 11 pe. Counterstaining with DAPI. Cross-section views created in Imaris Software show horizontal and vertical sections of enteroendocrine cells expressing HEV ORF2 antigens.(C) Presence of HEV capsid protein in enterocytes. Immunofluorescence staining of HEV ORF2 (green) and enterocytes (SI – sucrose isomaltase, red) in HEV-$3^{G1634R}$ electroporated HIEs at day 11 pe. Counterstaining with DAPI. (D) Representative images of mock electroporated fetal HIEs immunofluorescence staining at day 11 pe. Cells were stained with HEV ORF2-specific antibody (green) and antibody specific for proliferating cells (Ki67), enterocytes (SI, sucrose isomaltase), enteroendocrine cells (CHGA, chromogranin A) or goblet cells (WGA, wheat germ agglutinin). DAPI (blue) was used as counterstaining (nuclei). Scale bar – 50, 30 or 20 µm. pe, post-electroporation.

### Supplementary References

Author names in bold designate shared co-first authorship.
